# A Practical Alzheimer Disease Classifier via Brain Imaging-Based Deep Learning on 85,721 Samples

**DOI:** 10.1101/2020.08.18.256594

**Authors:** Bin Lu, Hui-Xian Li, Zhi-Kai Chang, Le Li, Ning-Xuan Chen, Zhi-Chen Zhu, Hui-Xia Zhou, Xue-Ying Li, Yu-Wei Wang, Shi-Xian Cui, Zhao-Yu Deng, Zhen Fan, Hong Yang, Xiao Chen, Paul M. Thompson, Francisco Xavier Castellanos, Chao-Gan Yan, the Alzheimer’s Disease Neuroimaging Initiative

**Affiliations:** CAS Key Laboratory of Behavioral Science, Institute of Psychology, Beijing, China; Department of Psychology, University of Chinese Academy of Sciences, Beijing, China; Center for Cognitive Science of Language, Beijing Language and Culture University, Beijing, China; Sino-Danish College, University of Chinese Academy of Science, Beijing, China; Sino-Danish Center for Education and Research, Beijing, China; Department of Neurosurgery, Huashan Hospital, Fudan University, Shanghai Neurosurgical Clinical Center, Shanghai, China; Department of Radiology, The First Affiliated Hospital, College of Medicine, Zhejiang University, Hangzhou, Zhejiang, China; Imaging Genetics Center, Mark & Mary Stevens Institute for Neuroimaging & Informatics, Keck School of Medicine, University of Southern California, Los Angeles, CA, USA; Department of Child and Adolescent Psychiatry, NYU Grossman School of Medicine, New York, NY, USA; Nathan Kline Institute for Psychiatric Research, Orangeburg, NY, USA; International Big-Data Center for Depression Research, Institute of Psychology, Chinese Academy of Sciences, Beijing, China; Magnetic Resonance Imaging Research Center, Institute of Psychology, Chinese Academy of Sciences, Beijing, China

**Keywords:** Alzheimer’s disease, convolutional neural network, magnetic resonance brain imaging, sex differences, transfer learning

## Abstract

Beyond detecting brain lesions or tumors, comparatively little success has been attained in identifying brain disorders such as Alzheimer’s disease (AD), based on magnetic resonance imaging (MRI). Many machine learning algorithms to detect AD have been trained using limited training data, meaning they often generalize poorly when applied to scans from previously unseen populations. Therefore, we built a practical brain MRI-based AD diagnostic classifier using deep learning/transfer learning on a dataset of unprecedented size and diversity. A retrospective MRI dataset pooled from more than 217 sites/scanners constituted one of the largest brain MRI samples to date (85,721 scans from 50,876 participants) between January 2017 and August 2021. Next, a state-of-the-art deep convolutional neural network, Inception-ResNet-V2, was built as a sex classifier with high generalization capability. The sex classifier achieved 94.9% accuracy and served as a base model in transfer learning for the objective diagnosis of AD. After transfer learning, the model fine-tuned for AD classification achieved 90.9% accuracy in leave-sites-out cross-validation on the Alzheimer’s Disease Neuroimaging Initiative (ADNI, 6,857 samples) dataset and 94.5%/93.6%/91.1% accuracy for direct tests on three unseen independent datasets (AIBL, 669 samples / MIRIAD, 644 samples / OASIS, 1,123 samples). When this AD classifier was tested on brain images from unseen mild cognitive impairment (MCI) patients, MCI patients who converted to AD were 3 times more likely to be predicted as AD than MCI patients who did not convert (65.2% vs 20.6%). Predicted scores from the AD classifier showed significant correlations with illness severity. In sum, the proposed AD classifier offers a medical-grade marker that has potential to be integrated into AD diagnostic practice.

## 1. Introduction

Magnetic resonance imaging (MRI) is widely used in neuroradiology to detect brain lesions including stroke, vascular disease, and tumor tissue. Still, MRI has been less useful in definitively identifying degenerative diseases including Alzheimer’s disease (AD), mainly because signatures of the disease are diffuse within the images and hard to distinguish from normal aging. Machine learning and deep learning methods have been trained on relatively small datasets, but limited training data often leads to poor generalization performance on new datasets not used the train the algorithms. In the current study, we aim to create a practical brain imaging-based AD classifier with high generalization capability via learning/transfer learning on a diverse range of large-scale datasets.

In recently updated AD diagnostic criteria, such as those proposed by International Working Group (IWG-2) Criteria for Alzheimer’s Disease Diagnosis and The National Institute on Aging – Alzheimer’s Association (NIA-AA) Alzheimer’s Diagnostic Framework, markers such as amyloid measures from cerebrospinal fluid (CSF) and amyloid-sensitive positron emission tomography (PET) have been integrated into the diagnosis of AD (1, 2). Diagnostic sensitivity and specificity have been greatly improved by these markers (1, 3). Even so, the invasive nature and lower availability of these markers limits their application in routine clinical settings. Thus, accurate diagnosis of AD and its early stage using a non-invasive and widely available technology is critically important. Structural MRI is a more promising candidate for imaging-based auxiliary diagnosis for AD considering its non-invasive nature and wider availability than PET. In addition, well-developed MRI data preprocessing pipelines make it feasible to integrate MRI markers into automatic end-to-end deep learning algorithms. Deep learning has already been successfully deployed in real-world scenarios such as extreme weather condition prediction (4), aftershock pattern prediction (5) and automatic speech recognition (6). In clinical scenarios, convolutional neural networks (CNN) – a widely-used architecture that is well-suited for image-based deep learning – have been successfully used for objective diagnosis of retinal diseases (7), skin cancer (8), and for breast cancer screening (9).

However, prior attempts at MRI-based AD diagnosis have yet to attain clinical utility. A major challenge for brain MRI-based algorithms, especially if they are trained on limited data, is their failure to generalize. For example, a brain imaging-based classifier may give precise predictions for testing samples from a specific hospital from which the training dataset came. However, performance of the classifier declines dramatically when directly applied to samples from another unknown hospital (10). One critical reason for performance discrepancy is that brain imaging data vary depending on scanner characteristics such as scanner vendor, magnetic field strength, head coil hardware, pulse sequence, applied gradient fields, reconstruction methods, scanning parameters, voxel size, field of view, etc. Participants also differ in sex, age, race/ethnicity, and education. Robust methods need to work well on diverse populations. These variations in the scans – and in the populations studied – make it hard for a brain imaging-based classifier trained on data from a single site (or a few sites) to generalize to data from unseen sites/scanners. This has prevented brain imaging-based classifiers from becoming practically useful in clinical settings. Most brain MRI-based studies either did not include independent validation (11-13) or did not achieve satisfactory performance in independent validations (14). In fact, reviews of brain imaging-based AD classifiers suggest that most machine learning methods have been trained on samples in the hundreds, with only 2 out of 81 studies (15), 0 out of 16 studies (16), and 6 out of 114 studies (17) (of those included in recent systematic reviews) including independent dataset validations, raising doubts about the generalizability of the models.

Another bottleneck in developing a practical brain imaging-based classifier involves the variety and comprehensiveness of training datasets. Directly training AD models on datasets that only contain several hundred samples may result in overfitting with poor generalization to unseen test data (15). The transfer learning framework has been proposed to solve this problem, by training a model on a certain characteristic for which abundant samples are available, and fine-tuning it to another characteristic, or for similar tasks, in smaller samples (18). Published evidence shows that pretrained models can outperform models trained from scratch in classification accuracy and robustness (19, 20). In medical imaging, transfer learning has been successfully applied to diagnose retinal disease (7) and skin cancer (8). Nonetheless, in brain imaging, no study has encompassed the tens of thousands of openly shared brain images to promote the generalizability of an AD classifier. Thus, in the current study, we used one of the largest and most diverse samples to date (N = 85,721 from more than 217 sites/scanners, see Table 1) to pre-train a brain imaging-based classifier with high generalizability. We chose a sex classifier rather than an age predictor as the base model for transfer learning, because age prediction error may contain biological meaning (e.g., increased predicted age may indicate accelerated aging (21)). Thus, it can be hard to measure the true performance of an age predictor while participant sex is more stable for classification. Subsequently, the pre-trained sex classifier was fine-tuned for AD classification and was validated through leave-sites-out cross-validation and three independent validations.

**Table 1:** Datasets used in the present study.

| Full Name of Dataset | Number of<br>T1 Scans | T1 Scans<br>after QC | Age<br>(mean±std) | Number of<br>Subjects | Number<br>of Sites | Manufacturers | Field<br>Strength |
| --- | --- | --- | --- | --- | --- | --- | --- |
| Adolescent Brain Cognition Development | 31176 | 30222 | 13.76±10.08 | 11875 | 21 | SIE/PHI/GE | 3T |
| UK Biobank | 20124 | 19744 | 63.1±7.46 | 20124 | 4 | SIE | 3T |
| Alzheimer's Disease Neuroimaging Initiative | 16596 | 16431 | 74.97±7.4 | 2546 | 57 | SIE/PHI/GE | 3T/1.5T |
| Open Access Series of Imaging Studies | 3150 | 3099 | 67.54±20.64 | 1664 | (5) | SIE | 3T/1.5T |
| REST-meta-MDD sample | 2380 | 2363 | 36.2±15.11 | 2380 | 17 | SIE/PHI/GE | 3T/1.5T |
| Brain Genomics Superstruct Project | 1570 | 1552 | 21.54±2.89 | 1570 | 2 | SIE | 3T |
| Human Connectome Project | 1267 | 1220 | / | 1267 | 1 | SIE | 7T/3T |
| Autism Brain Imaging Data Exchange | 1102 | 1073 | 17.09±8.06 | 1102 | 17 | SIE/PHI/GE | 3T/1.5T |
| Autism Brain Imaging Data Exchange II | 1043 | 1019 | 15.16±9.39 | 1043 | 19 | SIE/PHI/GE | 3T/1.5T |
| 1000 Functional Connectomes Project | 897 | 864 | 25.8±10.76 | 897 | 33 | SIE/PHI/GE | 3T/1.5T |
| ADHD-200 Sample | 876 | 864 | 12.35±3.28 | 876 | 8 | SIE/PHI | 3T/1.5T |
| Consortium for Reliability and Reproducibility | 714 | 691 | 23.45±12.31 | 715 | 2 | SIE/GE | 3T |
| Cambridge Centre for Ageing Neuroscience (Cam-CAN) | 652 | 523 | 54.36±18.55 | 652 | 1 | SIE | 3T |
| Enhanced Nathan Kline Institute - Rockland Sample | 646 | 616 | 38.63±21.21 | 646 | 1 | SIE | 3T |
| Southwest University Longitudinal Imaging Multimodal | 586 | 581 | 20.1±1.3 | 586 | 1 | SIE | 3T |
| Child Mind Institute Healthy Brain Network | 572 | 506 | 10.74±3.65 | 572 | 3 | SIE | 3T/1.5T |
| Establishing Moderators and Biosignatures of Antidepressant | 540 | 523 | / | 540 | 4 | SIE/PHI/GE | 3T |
| Southwest University Adult Lifespan Dataset | 493 | 483 | 45.16±17.45 | 493 | 1 | SIE | 3T |
| Max Planck Institute Leipzig Mind-Brain-Body Dataset | 316 | 291 | / | 316 | 1 | SIE | 3T |
| Beijing Enhanced Sample | 180 | 176 | 21.22±1.94 | 180 | 1 | SIE | 3T |
| Nathan Kline Institute - Rockland Sample | 167 | 151 | 35.59±20.71 | 167 | 1 | SIE | 3T |
| The Center for Biomedical Research Excellence | 147 | 137 | / | 147 | 1 | SIE | 3T |
| The Age-ility Project | 110 | 105 | 21.87±5.39 | 110 | 1 | SIE | 3T |
| Parkinson's Disease Datasets | 68 | 65 | 66.18±7.58 | 68 | 2 | SIE | 3T/1.5T |
| Power et al., 2012 Neuroimage Sample | 63 | 62 | 14.25±6.05 | 63 | 1 (2) | SIE | 3T |
| NYU Institute for Pediatric Neuroscience | 47 | 45 | 30.4±8.98 | 47 | 1 | SIE | 3T |
| Beijing Eyes Open Eyes Closed Sample | 46 | 44 | 22.54±2.18 | 46 | 1 | SIE | 3T |
| Multi-Modal MRI Reproducibility Resource | 42 | 42 | 31.76±9.35 | 42 | 1 | PHI | 3T |
| Adelstein et al., 2011, PLoS ONE Sample | 39 | 36 | 29.59±8.38 | 39 | 1 | SIE | 3T |
| Cleveland CCF | 31 | 29 | 43.55±11.14 | 31 | 1 | SIE | 3T |
| Virginia Tech Carilion Research Institute | 25 | 24 | 26.84±8.17 | 25 | 1 (3) | SIE | 3T |
| Beijing Short TR Sample | 24 | 23 | 23.71±6.74 | 24 | 1 | SIE | 3T |
| FIND Lab sample | 13 | 12 | 24.08±3.73 | 13 | 1 | GE | 3T |
| The Midnight Scan Club dataset | 10 | 10 | 29.1±3.35 | 10 | 1 | SIE | 3T |
|  | 85712 | 83735 |  | 50876 | 217 |  |  |

Note: Abbreviation: SIE = Siemens, Phi = Philips, GE = General Electric. Numbers in the bracket indicate the number of scanners.

The goal of the present study was to build a practical AD classifier with high generalizability. We incorporated three design features to improve the method’s clinical utility. First, we trained and tested the algorithm on a dataset of unprecedented size and diversity (from more than 217 sites/scanners). The variety of training samples was critical for improving model generalizability. Second, rigorous leave-datasets/sites-out cross-validation and independent validations were implemented to assure that classifier accuracy would be robust to site/scanner variability. Third, compared to 2D modules (feature detectors) typically used in CNNs for natural images, here, fully 3D convolution filters were used to capture more sophisticated and distributed spatial features for diagnostic classification. We also openly share our preprocessed data, trained model, code, and have built an online predicting website for anyone interested in testing our classifier.

## 2. Methods

### Data acquisition

We submitted data access applications to nearly all the open-access brain imaging data archives and received permissions from the administrators of 34 datasets. The full dataset list is shown in Table 1. Deidentified data were contributed from datasets collected with approvals from local Institutional Review Boards. The reanalysis of these data was approved by the Institutional Review Board of Institute of Psychology, Chinese Academy of Sciences. All participants had provided written informed consent at their local institution. All 50,876 participants (contributing 85,721 samples) had at least one session with a T1-weighted structural brain image and information on their sex and age. For participants with multiple sessions of structural images, each image was considered an independent sample for data augmentation in training. Importantly, scans from the same person were never split into training and testing sets, as that could artifactually inflate performance.

### MRI preprocessing

We did not feed raw data into the classifier for training but used accepted pre-processing pipelines that are known to generate useful features from brain scans. The brain structural data were segmented and normalized to acquire grey matter density (GMD) and grey matter volume (GMV) maps. Specifically, we used the voxel-based morphometry (VBM) analysis module within Data Processing Assistant for Resting-State fMRI (DPARSF) (22), which is based on SPM (23), to segment individual T1-weighted images into grey matter, white matter, and cerebrospinal fluid (CSF). Then, the segmented images were transformed from individual native space to MNI-152 space (a coordinate system created by Montreal Neurological Institute (24)) using the Diffeomorphic Anatomical Registration Through Exponentiated Lie algebra (DARTEL) tool (25). Two voxel-based structural metrics, GMD and GMV were fed into the deep learning classifier as two features for each participant. GMD is the output of the unmodulated tissue segmentation map in MNI space. GMV is calculated by multiplying the voxel value in GMD by the Jacobian determinants derived from the spatial normalization step (modulated) (26). Medical imaging-based classifiers could reach better or similar classification performances using an enhancing preprocessing procedure (27, 28).

### Quality control

Poor quality raw structural images would produce distorted GMD and GMV maps during segmentation and normalization. To remove such participants from affecting the training of the classifiers, we excluded participants in each dataset with a spatial correlation exceeding the threshold defined by (mean - 2SD) of the Pearson’s correlation between each participant’s GMV map and the grand mean GMV template. The grand mean GMV template was generated by randomly selecting 10 participants from each dataset (image quality visually checked for each participant) and averaging the GMV maps of all these 340 (from 34 datasets) participants. After quality control, 83,735 samples were retained for classifier training (Figure S1).

### Deep learning: classifier training and testing for sex

As the feature maps of brain MRI were three-dimensional (3D) rather than two-dimensional (2D), we could not directly use 2D pretrained models such as models trained based on ImageNet. Meanwhile, few pretrained 3D CNN models based on large-scale datasets, especially brain MRI datasets, exist. Therefore, we had to pretrain a brain MRI-based model for the further transfer learning procedure. It could have been more efficient to pretrain the model on another neurodegenerative disorder such as Parkinson’s disease (29, 30). However, we were unable to locate other neurodegenerative disorder datasets with tens of thousands of samples. Hence, we chose the sex classification task to pretrain the model because sex is a commonly available phenotype for any kind of datasets. We trained a 3-dimensional Inception-ResNet-v2(31) model adopted from its 2-dimensional version in the Keras built-in application (see Figure 1A for its structure). This is a state-of-the-art pattern recognition model, which integrates two classical series of CNN models, Inception and ResNet. We replaced the convolution, pooling, and normalization modules with their 3-dimensional versions and adjusted the number of layers and convolutional kernels to make them suitable for 3-dimensional MRI inputs (e.g., GMD and GMV as different input channels). The present model consists of one stem module, three groups of convolutional modules (Inception-ResNet-A/B/C) and two reduction modules (Reduction-A/B). The model can take advantage of convolutional kernels with different shapes and sizes, and can extract features of different sizes. The model also can mitigate vanishing gradients and exploding gradients by adding residual modules. We utilized the Keras built-in stochastic gradient descent optimizer with learning rate = 0.01, Nesterov momentum = 0.9, decay = 0.003 (e.g., learn rate = learn rate_0_ × (1 / (1 + decay × batch))). The loss function was set to binary cross-entropy. The batch size was set to 24 and the training procedure lasted 10 epochs for each fold. To avoid potential overfitting, we randomly split 600 samples out of the training sample as a validation sample and set a checking point at the end of every epoch. We saved the model in which the epoch classifier showed the lowest validation loss. Thereafter, the testing sample was fed into this model to test the classifier.

**Figure 1:**
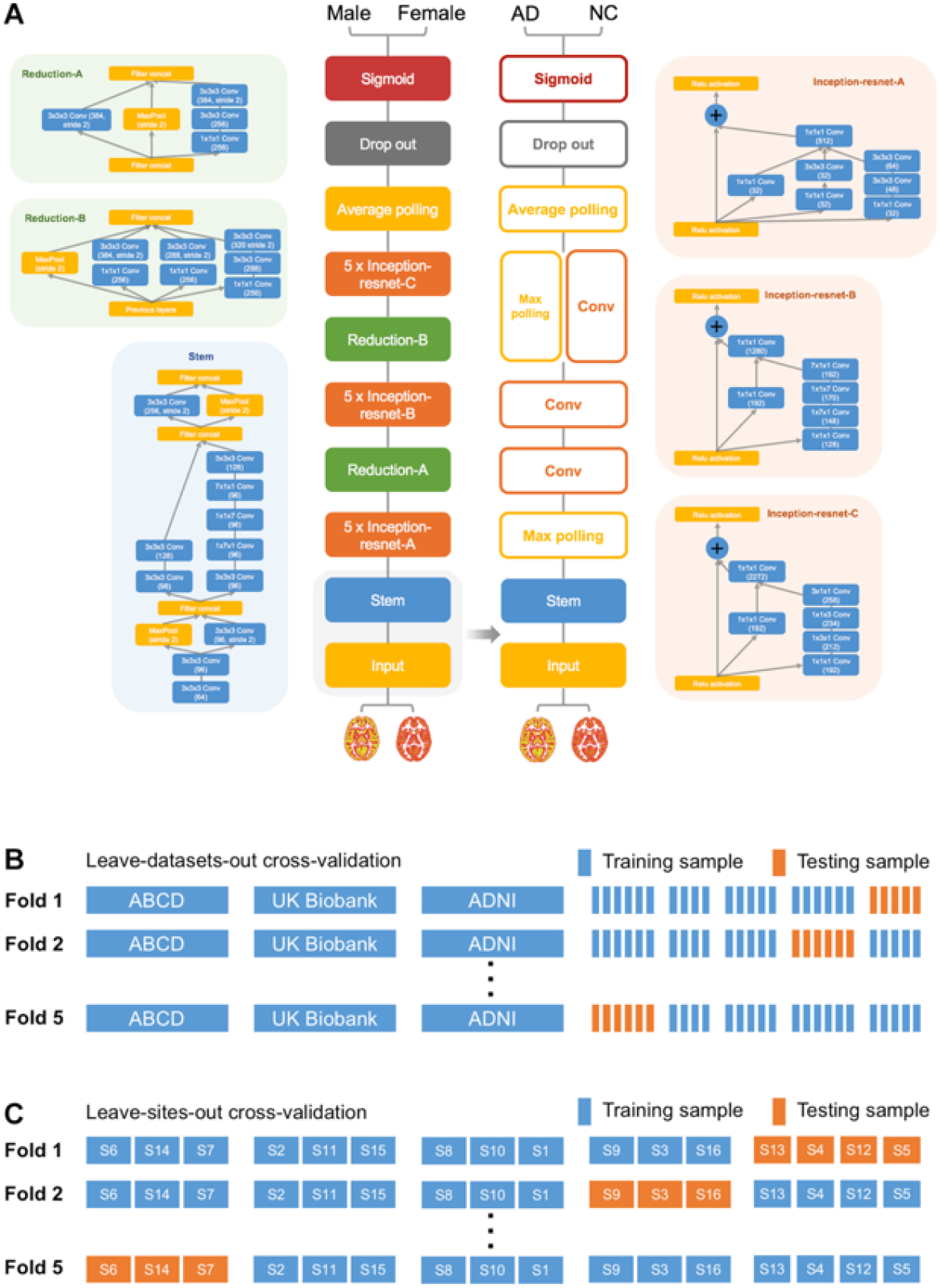
Flow diagram for the Alzheimer disease (AD) transfer learning framework and cross-validation procedure. (A) Schema for the 3D Inception-ResNet-V2 model and the transfer learning framework for the Alzheimer disease classifier. (B) Schematic diagram for the leave-datasets-out 5-fold cross-validation for the sex classifier. (C) Schematic diagram for the leave-sites-out 5-fold cross-validation for the AD classifier.

While training the sex classifier, random cross-validation may share participants from the same sites between training and testing samples, so the model may not generalize well to datasets from unseen sites due to site information leakage during training. To ensure generalizability, we used cross-dataset validation. In the testing phase, all the data from a given dataset would never be seen during the classifier training phase. This also ensured the data from a given site (and thus a given scanner) were unseen by the classifier during training (see Figure1B for an illustration). This strict setting can limit classifier performance, but it makes it feasible to generalize to any participant at any site (scanner). Five-fold cross-dataset validation was used to assess classifier accuracy. Of note, 3 datasets were always kept in the training sample due to the massive number of samples: Adolescent Brain Cognition Development (ABCD) (n = 31,176), UK Biobank (n = 20,124), and the Alzheimer’s Disease Neuroimaging Initiative (ADNI) (n = 16,596). The remaining 31 datasets were randomly allocated to training and testing samples. The allocating schemas were the solution that balanced the sample size of 5 folds the best from 10,000 random allocating procedures. Both healthy normal control and brain-related disorder patient samples in the 34 datasets were used to train the sex classifier.

### Transfer learning: classifier training and testing for AD

After obtaining a highly robust and accurate brain imaging-based sex classifier as a base model, we used transfer learning to further fine-tune the AD classifier. Rather than retaining the intact sophisticated structure of the base model (Inception-ResNet-V2), we only leveraged the pre-trained weights in the stem module and simplified the upper layers (e.g., replacing Inception-ResNet modules with ordinary convolutional layers). The retained bottom structure of the model works as a feature extractor and can take advantage of the massive training of the sex classifier. The pruned upper structure of the AD model can avoid potential overfitting and promote generalizability by reducing the number of parameters (10 million parameters for the AD classifier vs. 54 million parameters for the sex classifier). This derived AD classifier was fine-tuned on the ADNI dataset (2,186 samples from 380 AD patients and 4,671 samples from 698 normal controls (NCs), 76 ± 7 years, 3,493 samples from women). ADNI was launched in 2003 (Principal Investigator: Michael W. Weiner, MD) to investigate biological markers of the progression of MCI and early AD (see www.adni-info.org). We used the Keras built-in stochastic gradient descent optimizer with learning rate = 0.0003, Nesterov momentum = 0.9, decay = 0.002. The loss function was set to binary cross-entropy. The batch size was set to 24 and the training procedure lasted 10 epochs for each fold. Like the cross-dataset validation for sex classifier training, five-fold cross-site validation was used to assess classifier accuracy (see Figure 1C for an illustration). By ensuring that the data from a given site (and thus a given scanner) were unseen by the classifier during training, this strict strategy made the classifier generalizable with non-inflated accuracy, thus better simulating realistic clinical applications than traditional five-fold cross-validation. Other than using GMD+GMV as the input in transfer learning, we also used GMD, GMV or z-standardized normalized raw T1-weighted images as the input for the sex/AD classifiers to verify the influence of input format (Table 2). We also trained an age prediction model instead of the sex classifier in transfer learning to verify the influence of the base-model. We used the same structure as the sex classifier, except for adding a fully-connected layer with 128 neurons with “ELU” activation function before the final layer; we also changed the dropout rate from 0.5 to 0.2 following the parameters in the brain age prediction model reported by Jonsson et al. (21). Finally, we compared the performance of Inception-ResNet-V2 structure with the performances of some light-weight structures such as VGG19, DenseNet-201 and MobileNet-V2. The performances of these structures are listed in Table S1. In sum, the performance of models with light-weight structures were lower than that of the Inception-ResNet-V2 model, so that we chose Inception-ResNet-V2 for the model structure.

**Table 2:**
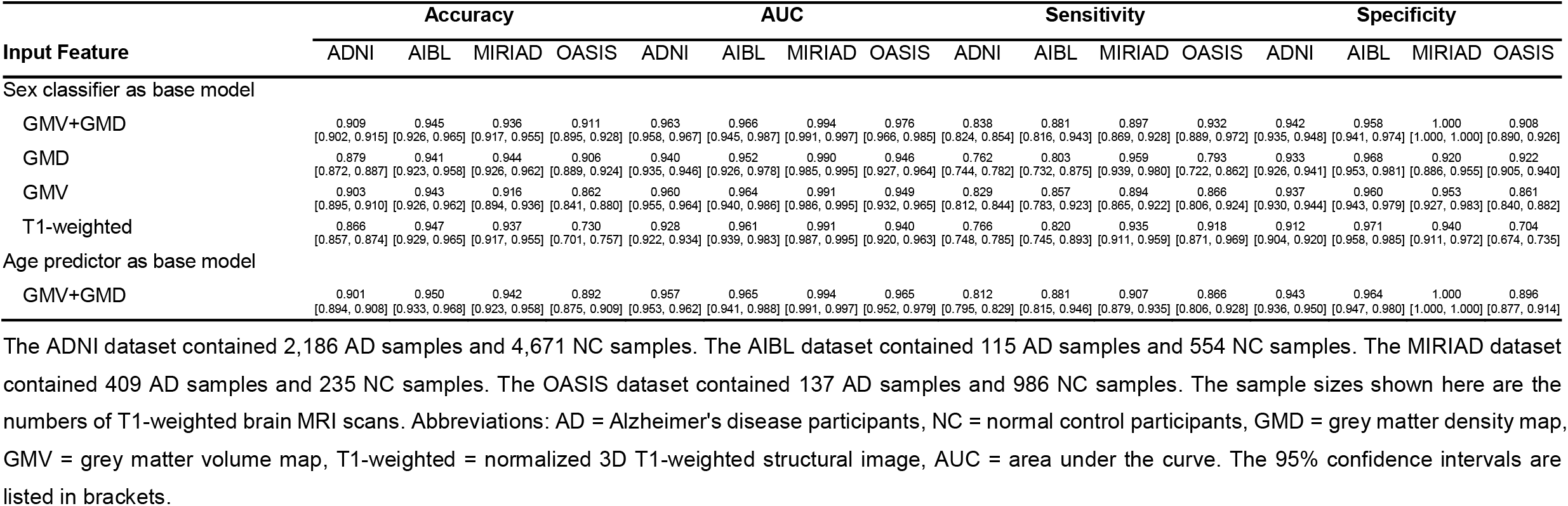
Performance of the Alzheimer’s disease classifier.

Furthermore, to test the generalizability of the AD classifier, we directly tested it on three unseen independent AD samples, i.e., the Australian Imaging, Marker and Lifestyle Flagship Study of Ageing (AIBL) (32), the Minimal Interval Resonance Imaging in Alzheimer’s Disease cohort (MIRIAD) (33), and the Open Access Series of Imaging Studies (OASIS) (34). We averaged the sigmoid activation output scores of the 5 AD classifiers in five-fold cross-validation on ADNI to obtain the final classification for each sample. We used diagnoses provided by the qualified physicians for the AIBL and MIRIAD datasets as the sample labels (115 samples from 82 AD patients and 554 samples from 324 NCs in AIBL, 74 ± 7 years, 374 samples from women; 409 samples from 46 AD patients and 235 samples from 23 NCs in MIRIAD, 70 ± 7 years, 358 samples from women). As OASIS did not specify the criteria for an AD diagnosis, we adopted criteria of MMSE and clinical dementia rating (CDR) modified from the ADNI-1 protocol manual to define AD and NC samples. Specifically, criteria for AD are (1) MMSE ≤ 22 and (2) CDR ≥ 1.0, and criteria for NC are (1) MMSE > 26 and (2) CDR = 0. Thus, we tested the model on 137 samples from 34 AD patients and 986 samples from 213 NC participants in the OASIS dataset after quality control, age 75 ± 10 years, 772 samples from women. Of note, the scanning conditions and recruitment criteria of these independent datasets differed much more than variations among different ADNI sites (where scanning and recruitment was deliberately coordinated), so we expected the AD classifier to achieve lower performance. We created heterogeneous distributions by randomly selecting 50% samples in each independent testing datasets 1,000 times to validate the stability of the model. The 95% confidence intervals of the classification performance metrics were produced from the random selection procedure.

We further investigated whether the AD classifier could predict disease progression in people with mild cognitive impairment (MCI). MCI is a syndrome defined as relative cognitive decline without symptoms interfering with daily life; even so, more than half of MCI patients progress to dementia within 5 years (35). The stable MCI (sMCI) samples were defined as “scans from an individual who was once diagnosed as MCI in any phase of ADNI and has not progressed to AD by the end of the ADNI follow-up”, and the progressive MCI (pMCI) samples were defined as “scans from a participant who was once diagnosed as MCI in any phase of ADNI and who has progressed to AD”. The scans labeled as “conversion” or “AD” (after conversion) for pMCI and the last scan for sMCI were excluded in the present study for precision. We screened imaging records of the MCI patients who converted to AD later in the ADNI 1/2/’GO’ phases, and collected 2,371 images from 243 participants labeled as ‘pMCI’. We also assembled 4,018 samples from 524 participants labeled ‘sMCI’ without later progression for contrast. We directly fed all these MCI images into the AD classifier without further fine-tuning, thus evaluating the performance of the AD classifier on unseen MCI information.

### Interpretation of the deep learning classifiers

To better understand the brain imaging-based deep learning classifier, we calculated occlusion maps for the classifiers. We repeatedly tested the images in the testing sample using the model with the highest accuracy within the 5 folds, while successively masking brain areas (volume = 18mm*18mm*18mm, step = 9mm) of all input images. The accuracy achieved on “intact” samples by the classifier minus accuracy achieved on “defective” samples indicated the “importance” of the occluded brain area for the classifier. The occlusion maps were calculated for both sex and AD classifiers. To investigate the clinical significance of the output of the AD classifier, we calculated the Spearman’s correlation coefficient between the predicted scores and MMSE scores of AD, NC, and MCI samples. We also used general linear models (GLM) to verify whether the predicted scores (or MMSE score) showed a group difference between people with sMCI and pMCI. The age and sex information of MCI participants was included in this GLM as covariates. We selected the T1-weighted images from the first visit for each MCI subject and finally collected data from 243 pMCI patients and 524 sMCI patients.

## 3. Results

### 3.1 Large-Scale Brain imaging Data

Only brain imaging data with enough size and variety can make deep learning accurate and robust enough to build a practical classifier. We received permissions from the administrators of 34 datasets (85,721 samples of 50,876 participants from more than 217 sites/scanners, see Table 1; some datasets did not require applications). After quality control, all these samples were used to pre-train the stem module to achieve better generalization for further AD classifier training. The T1-weighted images were collected through Magnetization-Prepared Rapid Gradient-Echo Imaging (MPRAGE) or Inversion Recovery Fast Spoiled Gradient Recalled Echo (IR-FSPGR) sequences of 1.5 tesla or 3 tesla MR scanners. The raw acquisition voxel sizes ranged from 0.7mm×0.7mm×0.7mm to 1.3mm×1.3mm×1.2mm. For further fine-tuning of the AD classifier, ADNI, AIBL, MIRIAD, and OASIS were selected to train and test the model.

### 3.2 Performance of the sex classifier

We trained a 3-dimensional Inception-ResNet-v2 model adapted from its 2-dimensional version in the Keras built-in application (see Figure 1A for structure). As noted in Methods, we did not feed raw data into the classifier for training, but used prior knowledge regarding helpful analytic pipelines. Grey matter density (GMD) and grey matter volume (GMV) maps were treated as different input channels for models. To ensure generalizability, five-fold cross-dataset validation was used to assess classifier accuracy. The five-fold cross-dataset validation accuracies were: 94.8%, 94.0%, 94.8%, 95.7%, and 95.8%. Taken together, accuracy was 94.9% in testing samples when pooling results across the five folds. The area under the curve (AUC) of the receiver operating characteristic (ROC) curve was the classifier performance index at various threshold settings (36, 37). The AUC of the sex classifier reached 0.981 (Figure 2). In short, our model can classify the sex of a participant based on brain structural imaging data from anyone on any scanner with an accuracy of about 95%. Interested readers can test this model on our online prediction website (http://brainimagenet.org).

**Figure 2:**
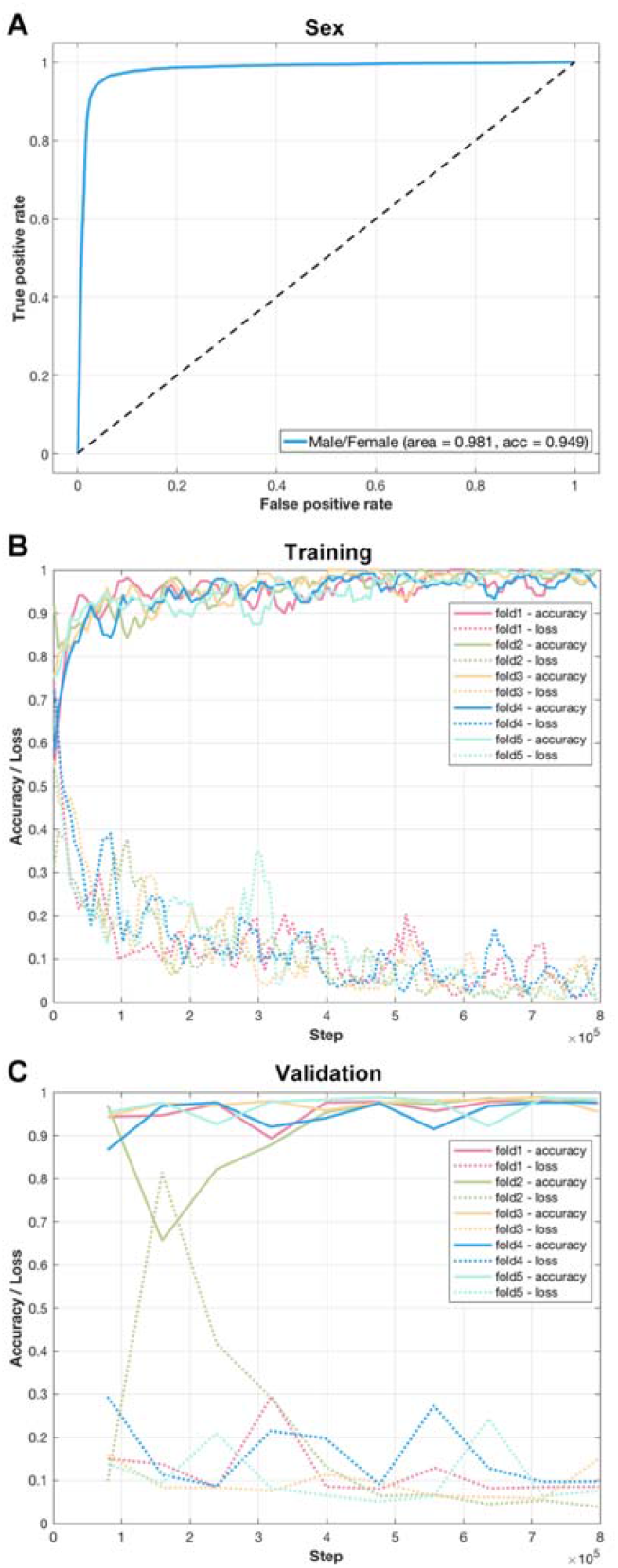
Performance of the sex classifier. (A) The receiver operating characteristic curve of the sex classifier. (B) The tensorboard monitor graph of the sex classifier in the training sample. The curve was smoothed for better visualization. (C) The tensorboard monitor graph of the sex classifier in the validation sample.

### 3.3 Performance of the AD classifier

After creating a practical brain imaging-based classifier for sex with high cross-dataset accuracy, we used transfer learning to see if we could classify patients with AD. The AD classifier achieved an accuracy of 90.9% (accuracy = 93.2%, 90.3%, 92.0%, 94.4%, and 86.7% in 5 cross-site folds) in the test samples. The 95% confidence interval of accuracy was [90.2%, 91.5%]. Average sensitivity and specificity were 0.838 and 0.942, respectively. The 95% confidence intervals of sensitivity and specificity were [0.824, 0.854] and [0.935, 0.948], respectively. The ROC AUC reached 0.963 when results from the 5 testing samples were taken together (see Figure 3 and Table 2). The 95% confidence interval of ROC AUC was [0.958, 0.967]. The AD classifier achieved an average accuracy of 91.4% on 3T field strength MR testing samples and achieved an average accuracy of 91.1% on 1.5T MR testing samples. The accuracy in 3T MR testing sample did not differ significantly from that of 1.5T MR testing sample (*p* = 0.316, by permutation test of randomly allocating the testing samples into 1.5T or 3T groups and calculating the accuracy difference between the two groups 100,000 times, Figure S2). In sum, the AUCs of models taking other types of images as the input (e.g., raw T1-weighted images) were slightly lower than that of the GMD+GMV image-based model. The GMD and GMV maps contained a priori knowledge from brain science, which might partly explain the better performance of GMD+GMV derived models.

**Figure 3:**
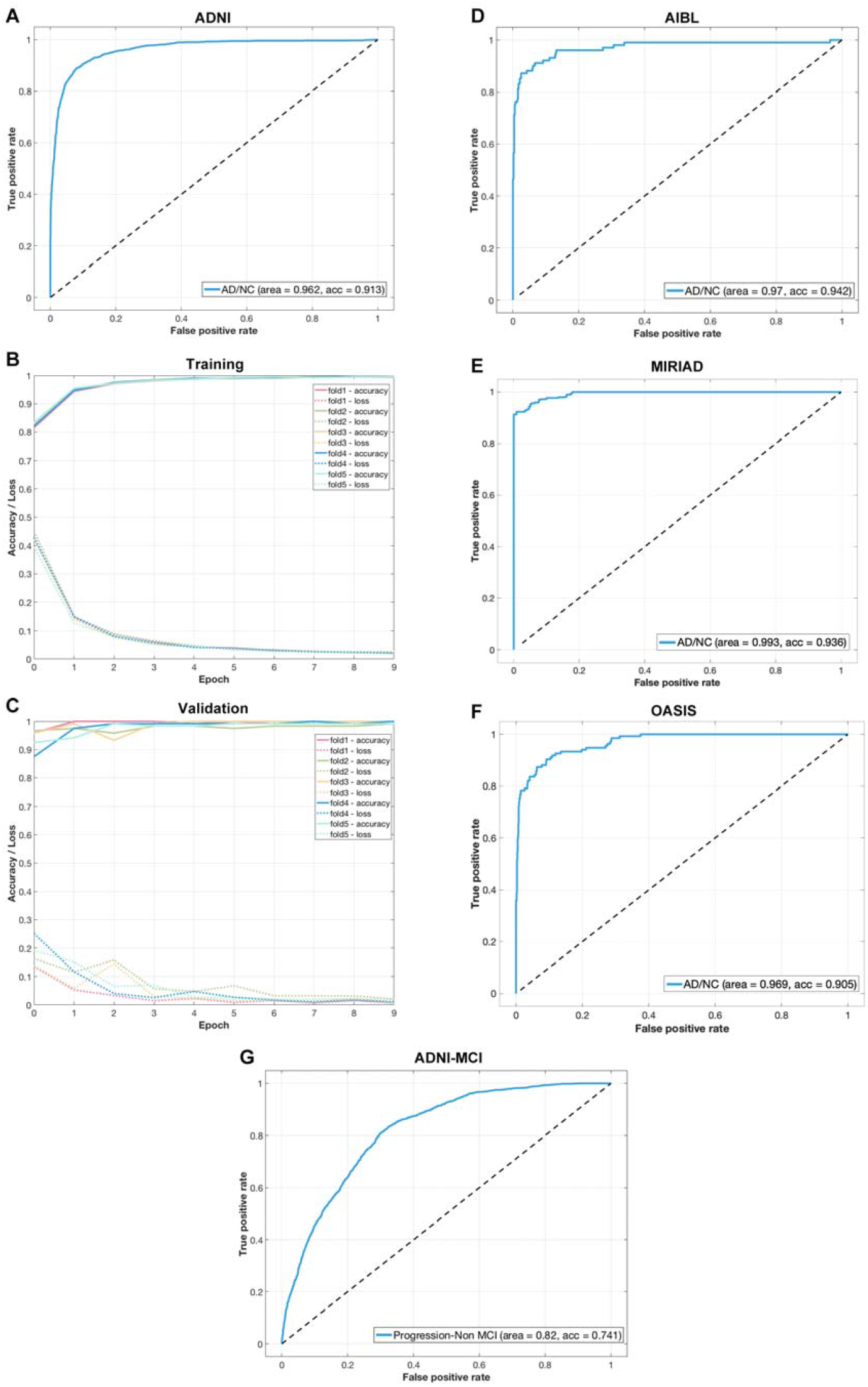
Performance of the Alzheimer’s disease (AD) classifier. Left panel shows the training and testing performance of the AD classifier on ADNI sample. Right panel shows the testing performance of the AD classifier on independent samples. (A) The receiver operating characteristic curve of the AD classifier. (B) The tensorboard monitor panel of the AD classifier in the training sample. (C) The tensorboard monitor panel of the AD classifier in the validation sample. (D) The ROC curve of AD classifier tested on the AIBL sample. (E) The ROC curve of the AD classifier tested on the MIRIAD sample. (F) The ROC curve of the AD classifier tested on the OASIS sample. (G) The ROC curve of the AD classifier tested on the ADNI MCI sample.

To test the generalizability of the AD classifier, we applied it to unseen independent AD datasets, i.e., AIBL, MIRIAD, and OASIS. The AD classifier achieved 94.5% accuracy in AIBL with 0.966 AUC (Figure 3D). Sensitivity and specificity were 0.881 and 0.958, respectively. The AD classifier achieved 93.6% accuracy in MIRIAD with 0.994 AUC (Figure 3E). Sensitivity and specificity were 0.897 and 1.000, respectively. The AD classifier achieved 91.1% accuracy in OASIS with 0.976 AUC (Figure 3F). Sensitivity and specificity were 0.932 and 0.908, respectively.

Importantly, although the AD classifier had never “seen” brain imaging data from subjects with MCI, we directly tested it on the MCI dataset in ADNI to see if it would have the potential to predict the progression of MCI to AD. We reasoned that even though people with MCI do not yet have AD, their scans may appear closer to the AD class learned by the deep learning model. We found that the AD classifier predicted 65.2% of pMCI patients as being in the AD class but only 20.4% of sMCI patients were predicted as having AD (Figure 3F). If the percentage of pMCI patients who were predicted as AD was considered as the sensitivity and the percentage of sMCI patients who were predicted as AD was considered as 1-specificity, the AUC of the ROC curve of the AD classifier reached 0.82. These results suggest that the classifier is practical for screening MCI patients to determine risk of progression to AD. In sum, we believe our AD classifier can provide important insights relevant to computer-aided diagnosis and prediction of AD, and we have freely provided it on the website http://brainimagenet.org. Importantly, classification results by the online classifier should be interpreted with caution, as they are probabilistic and cannot replace diagnosis by licensed clinicians.

As a supplementary analysis, we also tested transfer learning of the AD classifier using the intact structure of the base model (Figure S3). The performance of the model was uniformly somewhat inferior to the optimized AD classifier. The “intact” AD classifier achieved an average accuracy of 88.4% with 0.938 AUC in the ADNI test samples (Figure S4). Average sensitivity and specificity were 0.814 and 0.917, respectively. When tested on independent samples, the AD classifier achieved 91.2% accuracy in AIBL with 0.948 AUC. Sensitivity and specificity were 0.851 and 0.924, respectively. The AD classifier achieved 93.9% accuracy in MIRIAD with 0.995 AUC. Sensitivity and specificity were 0.905 and 0.996, respectively (Figure S5). The AD classifier achieved 86.1% accuracy in OASIS with 0.921 AUC. Sensitivity and specificity were 0.789 and 0.881, respectively. When tested on MCI samples, 63.2% of pMCI patients were predicted as having AD and 22.1% of sMCI patients were predicted as having AD by the AD classifier.

### 3.4 Interpretation of the deep learning classifiers

To better understand the brain imaging-based deep learning classifier, we calculated occlusion maps for the classifiers. The occlusion map showed that hypothalamus, superior vermis, pituitary, thalamus, amygdala, putamen, accumbens, hippocampus, and parahippocampal gyrus played critical roles in predicting sex (Figure 4A). The occlusion map for the AD classifier highlighted that the hippocampus and parahippocampal gyrus - especially in the left hemisphere - played unique roles in predicting AD (Figure 4B, Figure S6). Another visualization technique, Gradient-weighted Class Activation Mapping (Grad-CAM) (38), also showed relatively high weights of these regions in the input feature maps (Figure S9).

**Figure 4:**
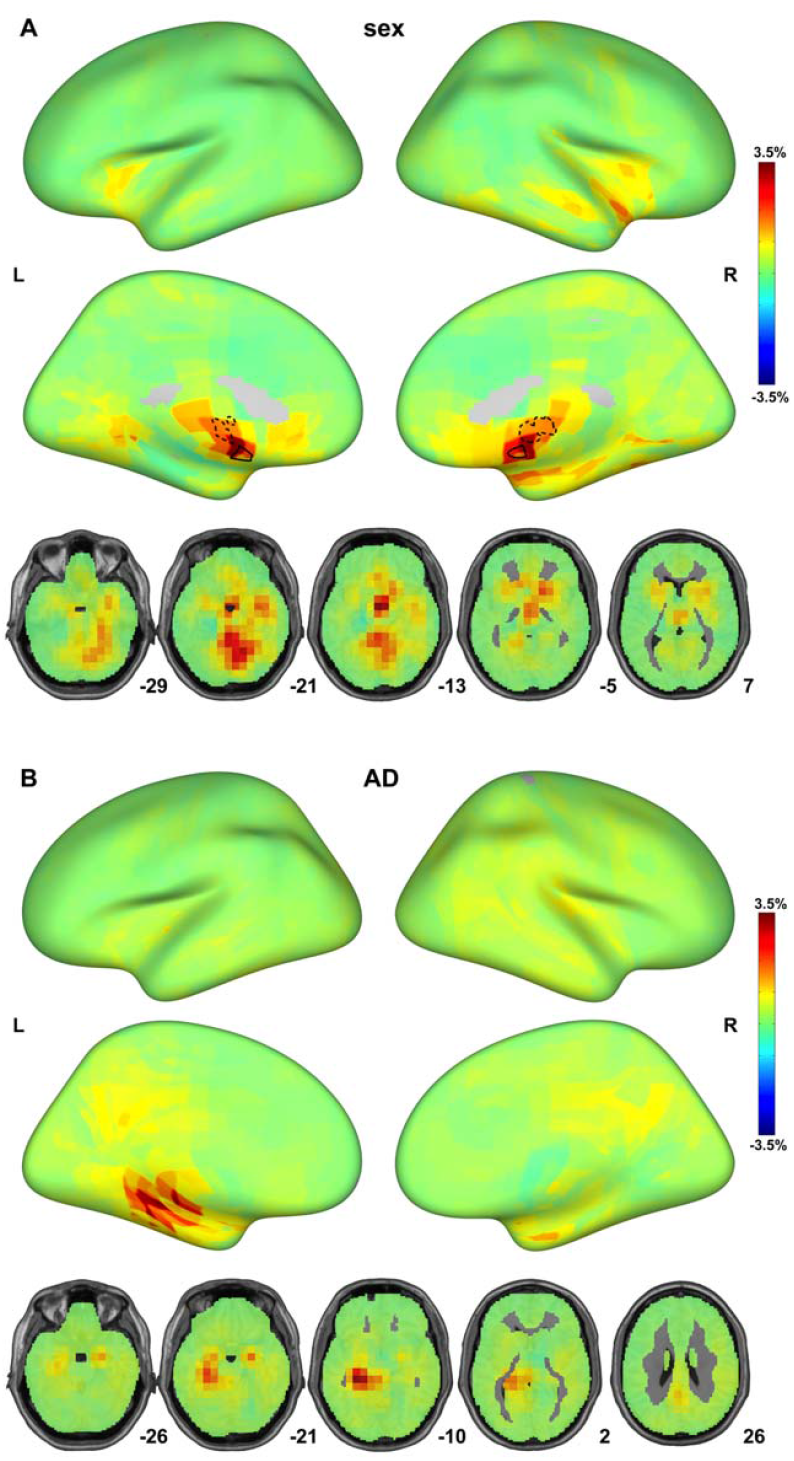
Interpretation of the deep learning classifiers with occlusion maps. Classifier performance dropped considerably when the brain areas rendered in red were masked out of the model input. (A) Occlusion maps for the sex classifier. Hypothalamus and pituitary were marked in dashed line and solid line. (B) Occlusion maps for the Alzheimer disease classifier.

To investigate the clinical significance of the output of the AD classifier, we calculated the Spearman’s correlation coefficient between the scores predicted by the classifier and MMSE scores in AD, NC, and MCI samples, although the classifier had not been trained with MMSE scores. This analysis confirmed significant negative correlations between the predicted scores and MMSE scores for AD (*r* = −0.37, *p* < 1 × 10^−55^), NC (*r* = −0.11, *p* < 1 × 10^−11^), MCI (*r* = −0.52, *p* < 1 × 10^−307^), and the overall samples (*r* = −0.64, *p* < 1 × 10^−307^) (Figure 5, Figure S7). As lower MMSE scores indicate more severe cognitive impairment in AD and MCI patients, we confirmed that the more severe the disease, the higher the classifier’s predicted score. In addition, both the predicted scores and MMSE scores differed significantly between pMCI and sMCI (predicted scores: *t* = 13.88, *p* < 0.001, Cohen’s d = 1.08; MMSE scores: *t* = −9.42, *p* < 0.01, Cohen’s d = −0.73, Figure S8). Importantly, the effect sizes of the classifier’s predicted scores were much larger than those for the behavioral measure (MMSE scores).

**Figure 5:**
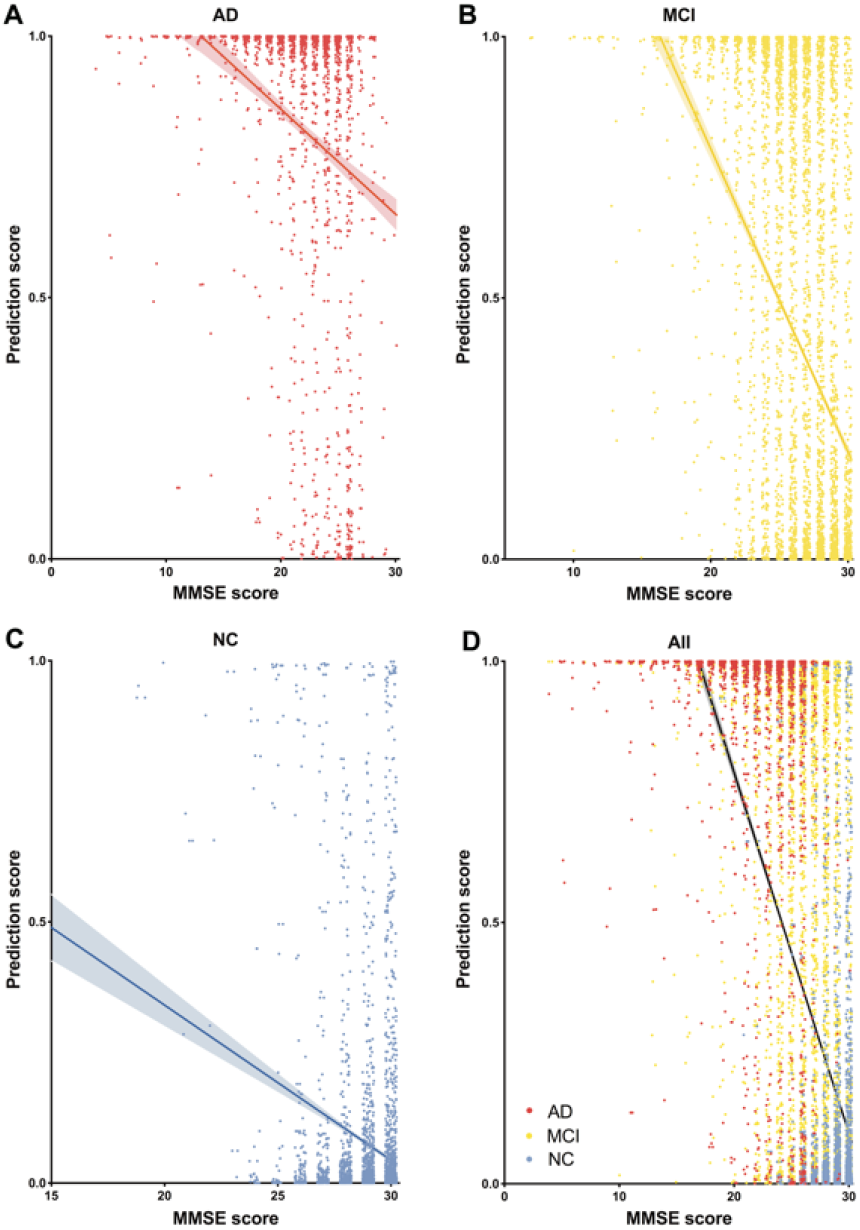
Correlations between the output of the Alzheimer’s disease (AD) classifier and severity of illness. The predicted scores from the AD classifier showed significant negative correlations with the mini-mental state examination (MMSE) scores of AD, normal control (NC) and mild cognitive impairment (MCI) samples. (A) Correlation between the predicted scores from the AD classifier and MMSE scores of AD samples. (B) Correlation between the predicted scores from the AD classifier and MMSE scores of MCI samples. (C) Correlation between the predicted scores from the AD classifier and MMSE scores of NC samples. (D) Correlation between the predicted scores from the AD classifier and the MMSE scores of AD, NC, and MCI samples, combined.

## 4. Discussion

Using an unprecedentedly diverse brain imaging sample, we pre-trained a sex classifier with about 95% accuracy which served as a base-model for transfer learning to promote model generalizability. After transfer learning, the model fine-tuned to AD achieved 90.9% accuracy in stringent leave-sites-out cross-validation and 94.2%/93.6%/91.1% accuracy for direct tests on three unseen independent datasets. Predicted scores from the AD classifier correlated significantly with illness severity. The AD classifier also showed the potential to predict the prognosis of MCI patients.

The high accuracy and generalizability of our deep neural network classifiers demonstrate that brain imaging has the practical potential to be auxiliary to the diagnostic process. One of the most prominent advantages of the present protocol is its outstanding generalizability, as validated by leave-sites-out validations and three independent-dataset validations. Performance of the AD classifier remained consistent despite considerable scanner/participant variations across four datasets used in the present study (e.g., ADNI, AIBL, MIRIAD and OASIS). Specifically, accuracies always exceeded 90% and AUCs always exceeded 0.96 in all four datasets. The present model outperformed models in recent studies whose accuracies range from 72.3% to 95% (39) or from 77% to 87% (14) using the same independent datasets (e.g., AIBL, MIRIAD, and OASIS). In addition, the analogous accuracies achieved on 1.5T and 3T MR ADNI imaging data further supported the robustness of the present classifier.

Of note, the output of the deep neural network model is a continuous variable, so the threshold can be adjusted to change sensitivity and specificity for certain purposes. For example, when tested on the AIBL dataset, sensitivity and specificity results were 0.881 and 0.958, respectively, as the default threshold was set at 0.5. However, for screening, the false-negative rate should be minimized even at the cost of higher false-positive rates. If we lower the threshold (e.g., to 0.2), sensitivity can be improved to 0.921 at a cost of decreasing specificity to 0.885. Thus, in our freely available AD prediction website, users can obtain continuous outputs and adjust the threshold to suit their specific purposes. Here, the sensitivity-specificity tradeoff of the AD classifier was consistent across different testing sites, so that physicians in diverse clinical settings can have consistent expectations for the classification tendencies of the classifier.

Beyond the feasibility of being integrated into the diagnostic process, the present AD model also showed potential to predict the progression of MCI patients. First, the present model was able to quantify key disease milestones by predicting disease progression in MCI patients. To wit, people with pMCI were 3 times more likely to be classified as AD than sMCI (65.2% vs 20.4%). Recently, a review on predicting progression from MCI noted that about 40% of studies had methodological issues, such as lack of a test dataset, data-leakage in feature selection or parameter tuning, and leave-one-out validation performance bias (40). The present AD classifier was only trained on AD/NC samples and was not fine-tuned using MCI data, so data leakage was avoided. The estimated true AUC of current published state-of-art classifiers for predicting progression of MCI is about 0.75 (15, 40). The proposed AD classifier here outperformed that benchmark (AUC = 0.82). Considering the discouraging clinical trial failures of AD treatments, early identification of people with MCI with high potential to progress to AD would help in the evaluation of early treatments (41).

Although deep-learning algorithms have often been referred to as “black boxes” for their poor interpretability, our subsequent analyses showed that the current MRI-based AD marker was aligned with pathological findings and clinical insights. For example, AD induced brain structural changes have been frequently reported by MRI studies. Among all the structural findings, hippocampal atrophy is the most prominent change and is used in imaging assisted diagnosis (42, 43). Neurobiological changes in the hippocampus typically precede progressive neocortical damage and AD symptoms. The convergence of our deep learning system and human physicians on alterations in hippocampal structure for classifying AD patients is in line with the crucial role of the hippocampus in AD. On the other hand, its maximum absolute value was only about 3.1%, which means that even if the most important brain area was eliminated from input, accuracy only dropped from 90.9% to about 88.8%. Interestingly, brain atrophy in AD has been frequently reported to be left lateralized (44, 45). Compared to the un-optimized AD classifier, a slight left hemisphere preference for input features may help explain the improved performance of the optimized AD classifier.

Rather than indiscriminately imitating the structure of the base model in transfer learning, the present AD classifier significantly simplified the model before the fine-tuning procedure. To wit, the performance of the unoptimized AD classifier was far poorer than that of the optimized AD classifier in accuracy, sensitivity, specificity, and in independent validation performance. Truncating or pruning models before transfer learning has been found to facilitate the performance of the transferred models (46, 47). As the sample for training the AD classifier is considerably smaller than that used to train the sex classifier, the simplified model structure may have helped to avoid overfitting and improve generalizability.

By precisely predicting sex, the present study provides evidence of sex differences in human brain. Daphna and colleagues extracted hundreds of VBM features from structural MRI and concluded that “the so-called male/female brain” does not exist as no individual structural feature supports a sexually dimorphic view of human brains (48), which was supported by a recent large-scale review (49). However, human brains may embody sexually dimorphic features in a multivariate manner. The high accuracy and generalizability of the present brain image-based sex classifier implies a “brain sex” is kind of recognizable in a 1,981,440-dimension (96*120*86*2) feature space, thus need to be further investigated in the future. Among those 1,981,440 features, the hypothalamus and pituitary played the most critical roles in predicting sex. The hypothalamus regulates testosterone secretion through the hypothalamic-pituitary-gonadal axis and thus plays a critical role in brain masculinization (50). Men have significantly larger hypothalamus than women relative to cerebrum size (51). In addition, cerebellum – especially the vermis – strongly contributed to sex classification, in line with MRI morphology studies of sex differences in cerebellum (52, 53). Taken together, our machine learning evidence suggests that male/female brain differences do exist, in the sense that accurate classification is possible.

In the deep learning field, the emergence of ImageNet tremendously accelerated the evolution of computer vision (54). ImageNet provided large amounts of well-labeled image data for researchers to pre-train their models. Studies have shown that pre-trained models can facilitate the performance and robustness of subsequently fine-tuned models (19). The present study confirms that the “pre-train + fine-tuning” paradigm also works for MRI-based auxiliary diagnosis. Unfortunately, no such a well-preprocessed dataset exists in the brain imaging domain. As data organization and preprocessing of MRI data require tremendous time, manpower and computational load, these constraints impede scientists from other fields entering brain imaging. Open access to large amounts of preprocessed brain imaging data is fundamental to facilitate the participation of a broader range of researchers. Beyond building and sharing a practical brain imaging-based deep learning classifier, we will openly share all sharable preprocessed data to invite researchers (especially computer scientists) to join the efforts to create predictive models using brain images (Link_To_Be_Added upon publication; preprocessed data of some datasets will not be shared as the raw data owners do not allow sharing of data derivatives). We anticipate that this dataset may boost the clinical utility of brain imaging as ImageNet has done in computer vision research. We openly share our models to allow other researchers to deploy them (https://github.com/Chaogan-Yan/BrainImageNet). Our code is also openly shared as well, allowing other researchers to replicate the present results and further develop brain imaging-based classifiers based on our existing work. Finally, we have also built a demonstration website for classifying sex and AD (http://brainimagenet.org). Users can upload raw T1-weighted or preprocessed GMD and GMV data to make predictions of sex or AD labels in real-time.

Limitations of the current study should be acknowledged. Considering the lower reproducibility of functional MRI compared to structural MRI, only structural MRI derived images were used in the present deep learning model. Even so, functional measures of physiology and activation may further improve the performance of sex and brain disorder classifiers. In future studies, functional MRI, especially resting-state functional MRI, may provide additional information for model training. Furthermore, with advances in software such as FreeSurfer(55), fmriprep(56), and DPABISurf, surface-based algorithms have shown their superiority when compared with traditional volume-based algorithms (57). Surface-based algorithms are more time consuming to run in terms of computation load, but can provide more precise brain registration and reproducibility. Future studies should take surface-based images as inputs for deep learning models. In addition, the present AD classification model was built based on labels provided by the ADNI database. Using post-mortem neuropathological data, the gold standard for AD diagnoses, could further advance the clinical value of MRI-based markers.

In summary, we pooled MRI data from more than 217 sites/scanners to constitute one of the largest brain MRI samples to date, with the preprocessed imaging data derivatives openly shared to the scientific community whenever allowed. The brain imaging-based AD classifier derived from transfer learning achieved both high rates of accuracy and generalizability, which were validated by strict cross-sites-validation and independent datasets validation. The AD classifier was able to predict the progression of MCI patients non-invasively. The present study demonstrates the feasibility of the transfer learning framework in brain disorder applications. Future work should examine such a framework to assess psychiatric disorders, to predict treatment response, and individual differences more broadly.

## Declarations

## Ethics approval and consent to participate

Deidentified data were contributed from datasets collected with approval by local Institutional Review Boards. The reanalysis of these data was approved by the Institutional Review Board of Institute of Psychology, Chinese Academy of Sciences. All participants had provided written informed consent at their local institutions.

## Consent for publication

Not applicable.

## Availability of data and materials

The imaging, phenotype and clinical data used for the training, validation and test sets were obtained by application from the administrators of 34 datasets. The preprocessed brain imaging data for which raw data owners permit sharing data derivatives will be available on (Link_To_Be_Added upon publication). The code for training and testing the model are openly shared at https://github.com/Chaogan-Yan/BrainImageNet. Demonstration website for classifying sex and AD is available at http://brainimagenet.org.

## Authors’ contributions

C.-G.Y. designed the overall experiment. B.L., H.-X.L., L.L., N.-X.C., Z.-C.Z., H.-X.Z., X.-Y.L., Y.-W.W., S.-X.C., Z.-Y.D., Z.F., H.Y. and X.C. applied and preprocessed imaging data. H.-X.L. and B.L sorted the phenotype information of datasets. B.L. designed the model architectures and trained the models, B.L., Z.-K.C and C.-G.Y. built the online classifiers. C.-G.Y. provided technical supports and supervised the project. B.L. and C.-G.Y. wrote the paper, P.M.T edited the paper and suggested supplementary analysis, F.X.C. edited and polished the paper. All authors approved the manuscript and had final responsibility for the decision to submit for publication.

## Acknowledgements

Data used in the preparation of this article for training and testing the sex classifier was obtained from the Adolescent Brain Cognitive Development (ABCD) Study (https://abcdstudy.org), held in the NIMH Data Archive (NDA). This is a multisite, longitudinal study designed to recruit more than 10,000 children ages 9-10 and follow them over 10 years into early adulthood. The ABCD Study is supported by the National Institutes of Health and additional federal partners under award numbers U01DA041048, U01DA050989, U01DA051016, U01DA041022, U01DA051018, U01DA051037, U01DA050987, U01DA041174, U01DA041106, U01DA041117, U01DA041028, U01DA041134, U01DA050988, U01DA051039, U01DA041156, U01DA041025, U01DA041120, U01DA051038, U01DA041148, U01DA041093, U01DA041089. A full list of supporters is available at https://abcdstudy.org/federal-partners.html. A listing of participating sites and a complete listing of the study investigators can be found at https://abcdstudy.org/scientists/workgroups/. ABCD consortium investigators designed and implemented the study and/or provided data but did not necessarily participate in analysis or writing of this report. This manuscript reflects the views of the authors and may not reflect the opinions or views of the NIH or ABCD consortium investigators. This research has been conducted using the UK Biobank Resource. Data collection and sharing for the training and testing the sex and AD classifiers were funded by the Alzheimer’s Disease Neuroimaging Initiative (ADNI) (National Institutes of Health Grant U01 AG024904) and DOD ADNI (Department of Defense award number W81XWH-12-2-0012). ADNI is funded by the National Institute on Aging, the National Institute of Biomedical Imaging and Bioengineering, and through generous contributions from the following: AbbVie; Alzheimer’s Association; Alzheimer’s Drug Discovery Foundation; Araclon Biotech; BioClinica, Inc.; Biogen; Bristol-Myers Squibb Company; CereSpir, Inc.; Cogstate; Eisai Inc.; Elan Pharmaceuticals, Inc.; Eli Lilly and Company; EuroImmun; F. Hoffmann-La Roche Ltd and its affiliated company Genentech, Inc.; Fujirebio; GE Healthcare; IXICO Ltd.; Janssen Alzheimer Immunotherapy Research & Development, LLC.; Johnson & Johnson Pharmaceutical Research & Development LLC.; Lumosity; Lundbeck; Merck & Co., Inc.; Meso Scale Diagnostics, LLC.; NeuroRx Research; Neurotrack Technologies; Novartis Pharmaceuticals Corporation; Pfizer Inc.; Piramal Imaging; Servier; Takeda Pharmaceutical Company; and Transition Therapeutics. The Canadian Institutes of Health Research is providing funds to support ADNI clinical sites in Canada. Private sector contributions are facilitated by the Foundation for the National Institutes of Health (www.fnih.org). The grantee organization is the Northern California Institute for Research and Education, and the study is coordinated by the Alzheimer’s Therapeutic Research Institute at the University of Southern California. ADNI data are disseminated by the Laboratory for Neuro Imaging at the University of Southern California. Data used in the preparation of this article were obtained from the MIRIAD database. The MIRIAD investigators did not participate in analysis or writing of this report. The MIRIAD dataset is made available through the support of the UK Alzheimer’s Society (Grant RF116). The original data collection was funded through an unrestricted educational grant from GlaxoSmithKline (Grant 6GKC).

## Authors’ information

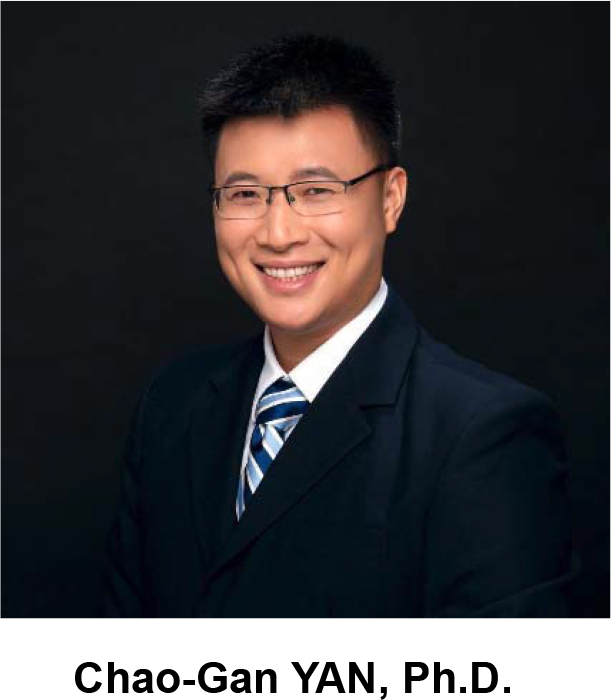

Dr. Chao-Gan Yan is a professor at the Institute of Psychology, Chinese Academy of Sciences (IPCAS). He is the Director of the Magnetic Resonance Imaging Research Center, the Director of the International Big-Data Center for Depression Research, and the Principal Investigator of The R-fMRI Lab located at IPCAS. His work has been widely cited in the scientific community (total citations > 16000, http://scholar.google.com/citations?user=lJQ9B58AAAAJ), achieving an h-index of 38. Six of his first-author/corresponding author papers ranked as ESI Top 1% Highly Cited Papers, 2 of which were ESI Top 1‰. He has been ranked as 2019, 2020 and 2021 most cited Chinese Researchers by Elsevier. He was awarded the 2021 Early Career Investigator Award by The Organization for Human Brain Mapping (OHBM). Additionally, he currently serves as associate editor for Neuroimage: Reports and serves on the editorial boards of NeuroImage and Journal of Neuroscience Methods. His research mainly focuses on resting-state fMRI (R-fMRI) computational methodology, mechanisms of spontaneous brain activity, and their applications in depression. He has addressed fundamental methodological issues (e.g., impact of head motion (ESI Top 1‰ of highly cited papers), standardization (ESI Top 1% of highly cited papers) and multiple comparison correction (ESI Top 1% of highly cited papers)) on the study of resting-state functional connectomics. He has also developed data processing and analysis toolbox for R-fMRI, DPABI (ESI Top 1‰ of highly cited papers) and DPARSF, which has been cited over 2000 times. He initiated a Chinese consortium for big data of brain imaging of depression (DIRECT), performed a big data study of depression neuroimaging (PNAS 2019, ESI Top 1% of highly cited papers), and studied the brain mechanisms of depression through a longitudinal study of an animal model. He has published 70+ peer-reviewed articles (30+ as first or corresponding author) in prestigious journals including PNAS, Molecular Psychiatry, NeuroImage, Human Brain Mapping.

