## Supplemental Materials for "A Practical Alzheimer Disease Classifier via Brain Imaging-Based Deep Learning on 85,721 Samples"

**Supplementary Materials**

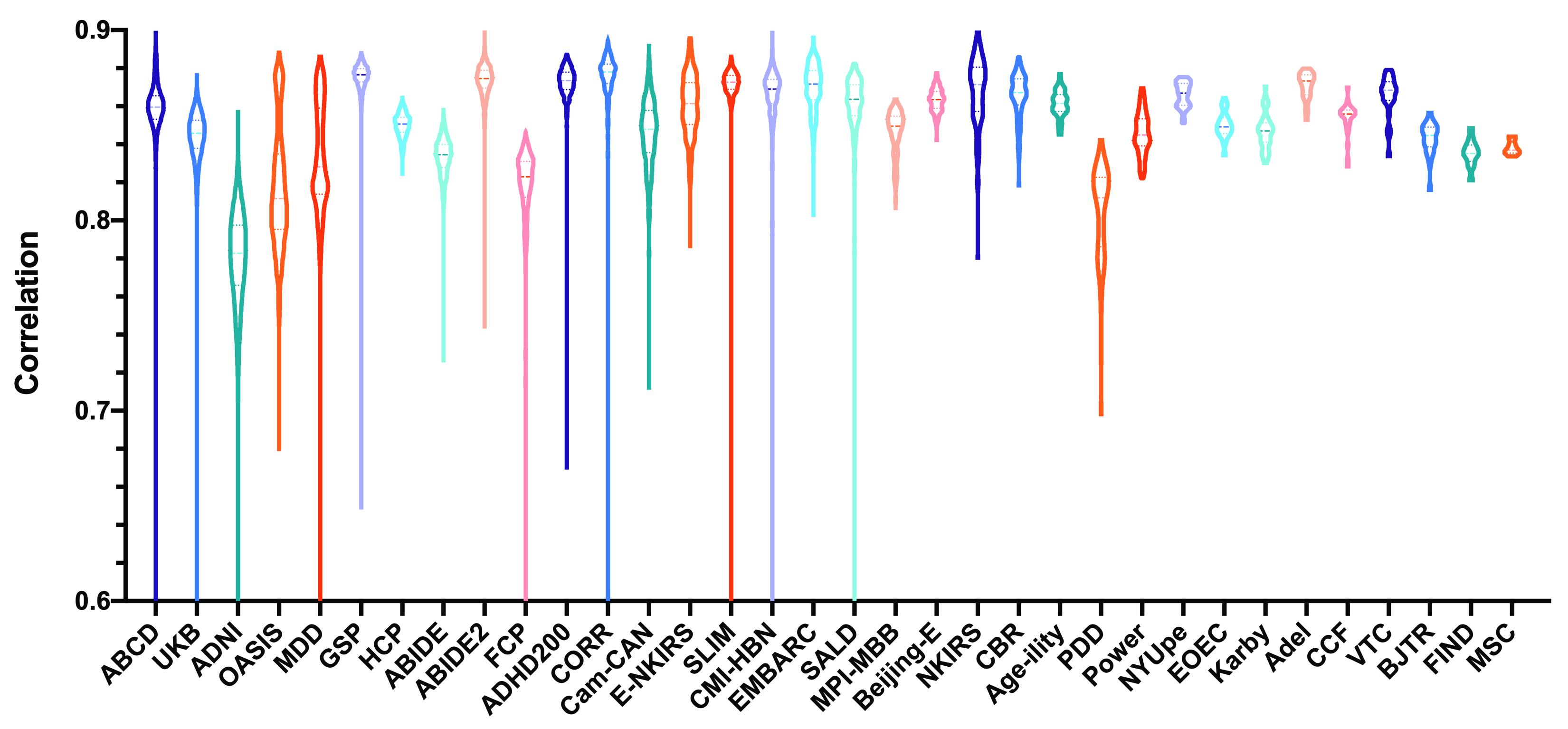

***Figure S1:* Violin plot of the distribution of correlations between grand-mean grey matter volume (GMV) template and GMV samples in each dataset**

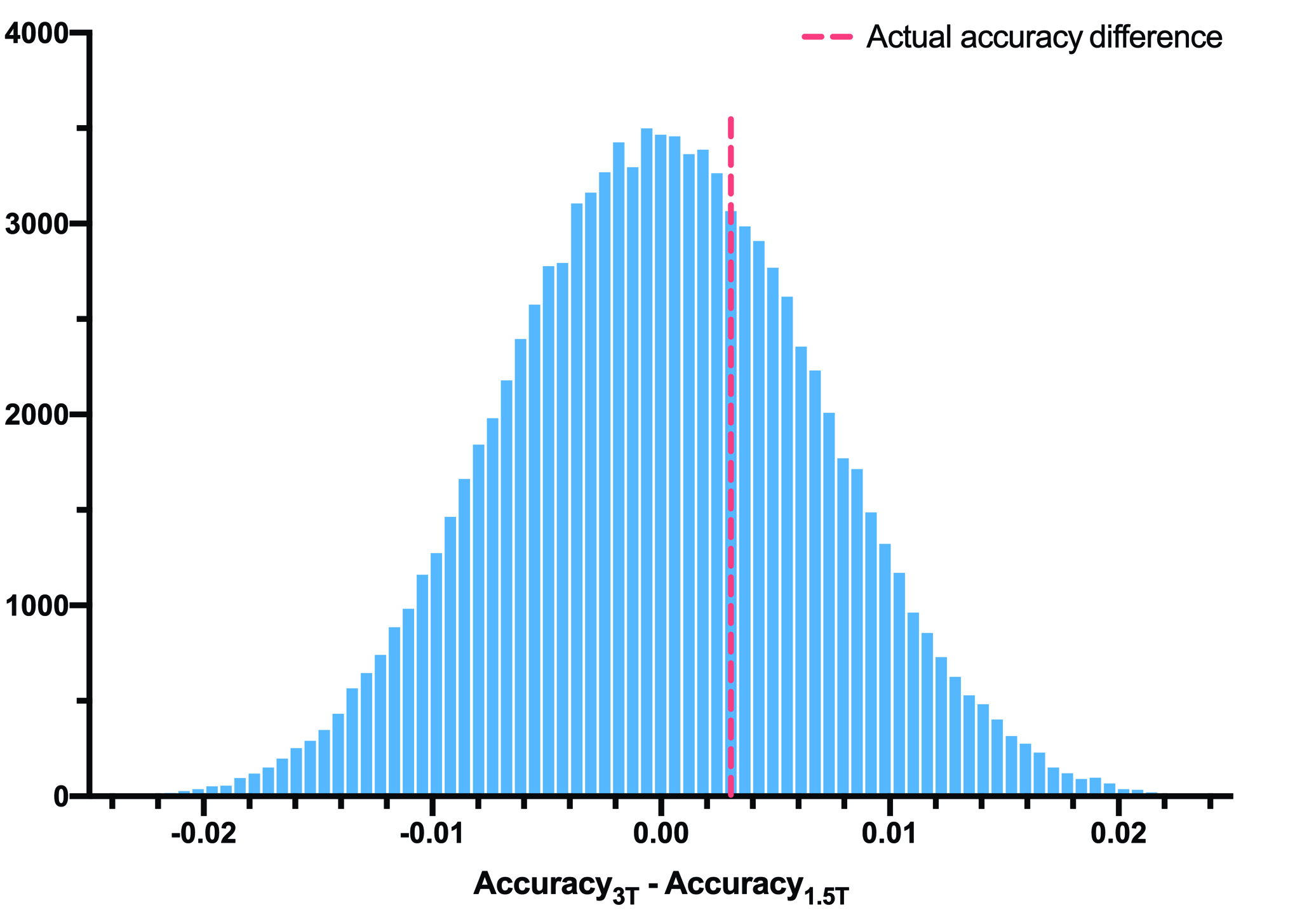

***Figure S2:* The distribution of 100,000 accuracy differences between randomly permutated pseudo 3T samples and pseudo 1.5T samples**

Accuracy on 3T-MR ADNI samples was 91.4% and accuracy on 1.5T-MR ADNI samples was 91.1% in five-fold cross-validation. The actual “3T vs 1.5T accuracy difference” was 91.4%-91.1%=0.3%, which is denoted by the red dashed line. Permutation testing was used to test whether the actual “3T vs 1.5T accuracy difference” (0.3%) differed significantly from zero. Specifically, we randomly allocated 4,068 samples into the “pseudo 3T group” and allocated 2,724 samples into the “pseudo 1.5T group”. The prediction accuracies of the two pseudo groups were computed and the accuracy difference in a random instance was obtained. The random allocation was repeated 10,000 times yielding a distribution of the randomized accuracy differences. The “actual accuracy difference” (0.3%) was located within the 95% confidence interval of this distribution.

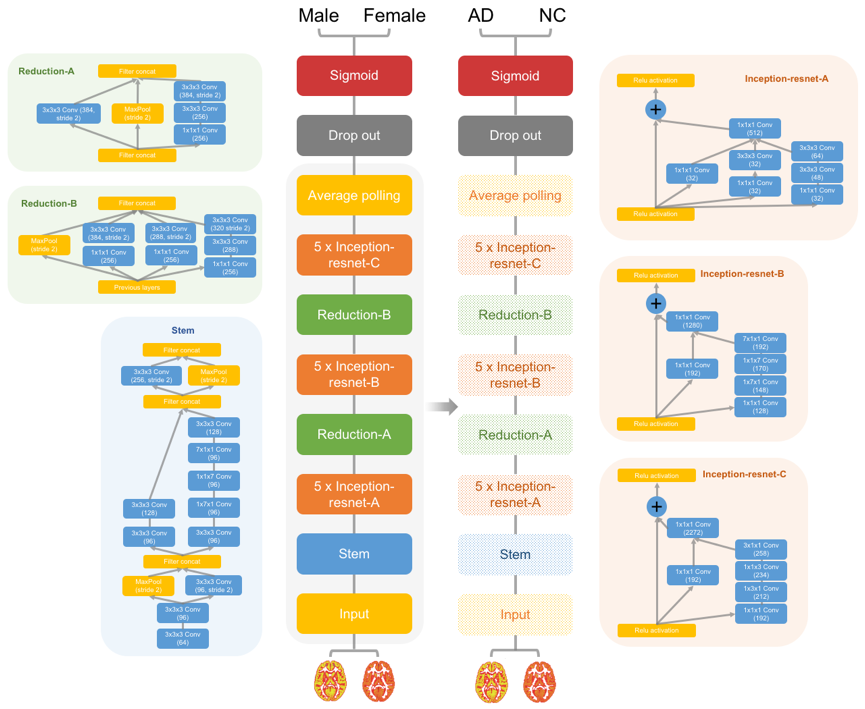

***Figure S3:* Flow diagram for the Alzheimer disease transfer learning framework without optimization**

Schema for 3D Inception-ResNet-V2 model and the Alzheimer disease classifier without model structure optimizing before transfer learning.

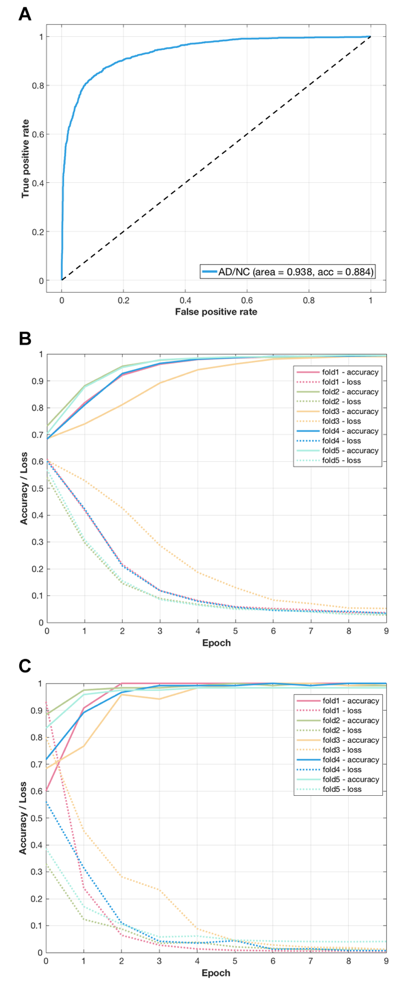

***Figure S4:* Performance of the Alzheimer's disease (AD) classifier without optimization**

(A) The receiver operating characteristic curve of the AD classifier. (B) The tensorboard monitor panel of the AD classifier in the training sample. (C) The tensorboard monitor panel of the AD classifier in the validation sample.

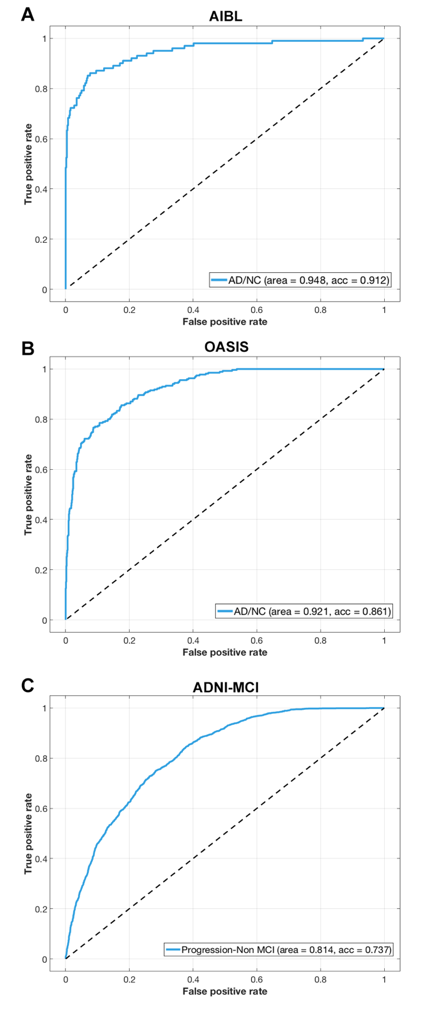

***Figure S5:* Receiver operating characteristic curves of the Alzheimer disease (AD) classifier without optimization when tested on independent AD samples and a mild cognitive impairment sample**

(A) The ROC curve of AD classifier tested on the AIBL sample. (B) The ROC curve of AD classifier tested on the OASIS sample. (C) The ROC curve of AD classifier tested on MCI sample in ADNI. The images of MCI subjects with future conversion to AD were labeled as “AD”, and the images of MCI subjects which hadn’t shown conversion to AD were labeled as “NC”.
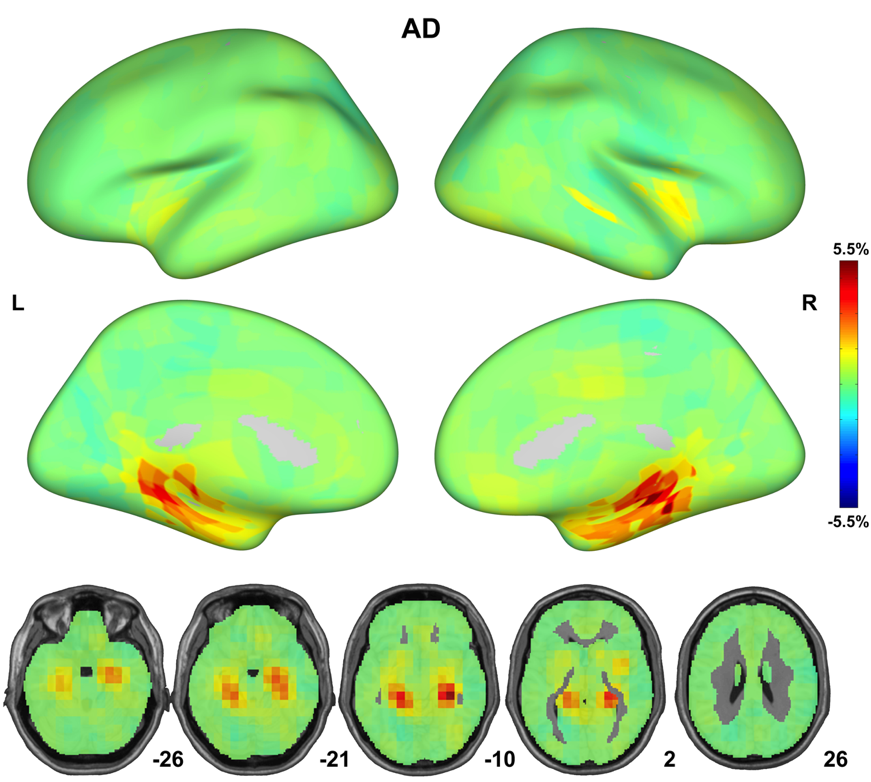

***Figure S6:* Interpretation of the AD classifiers without optimization by occlusion maps**

Classifier performance dropped considerably when the brain areas rendered in red were masked out of the model input.

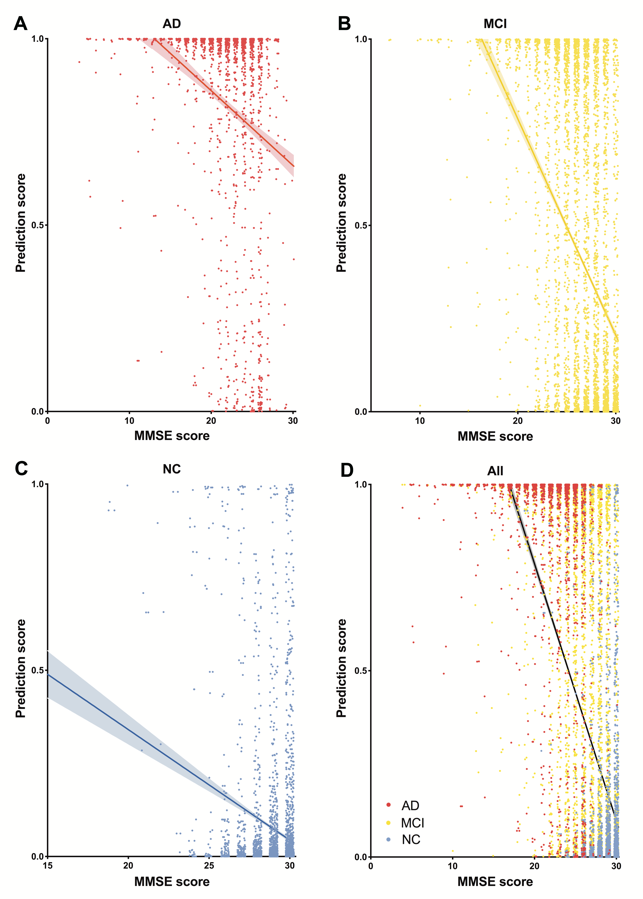

***Figure S7:*** **Correlations between the output of the Alzheimer's disease (AD) classifier without optimization and the severity of illness**

The predicted scores from the AD classifier showed significant negative correlations with the mini-mental state examination (MMSE) scores of AD, normal control (NC) and mild cognitive impairment (MCI) samples. (A) Correlation between the predicted scores from the AD classifier and MMSE scores of AD samples. (B) Correlation between the predicted scores from the AD classifier and MMSE scores of MCI samples. (C) Correlation between the predicted scores from the AD classifier and MMSE scores of NC samples. (D) Correlation between the predicted scores from the AD classifier and combined MMSE scores of AD, NC, and MCI samples.

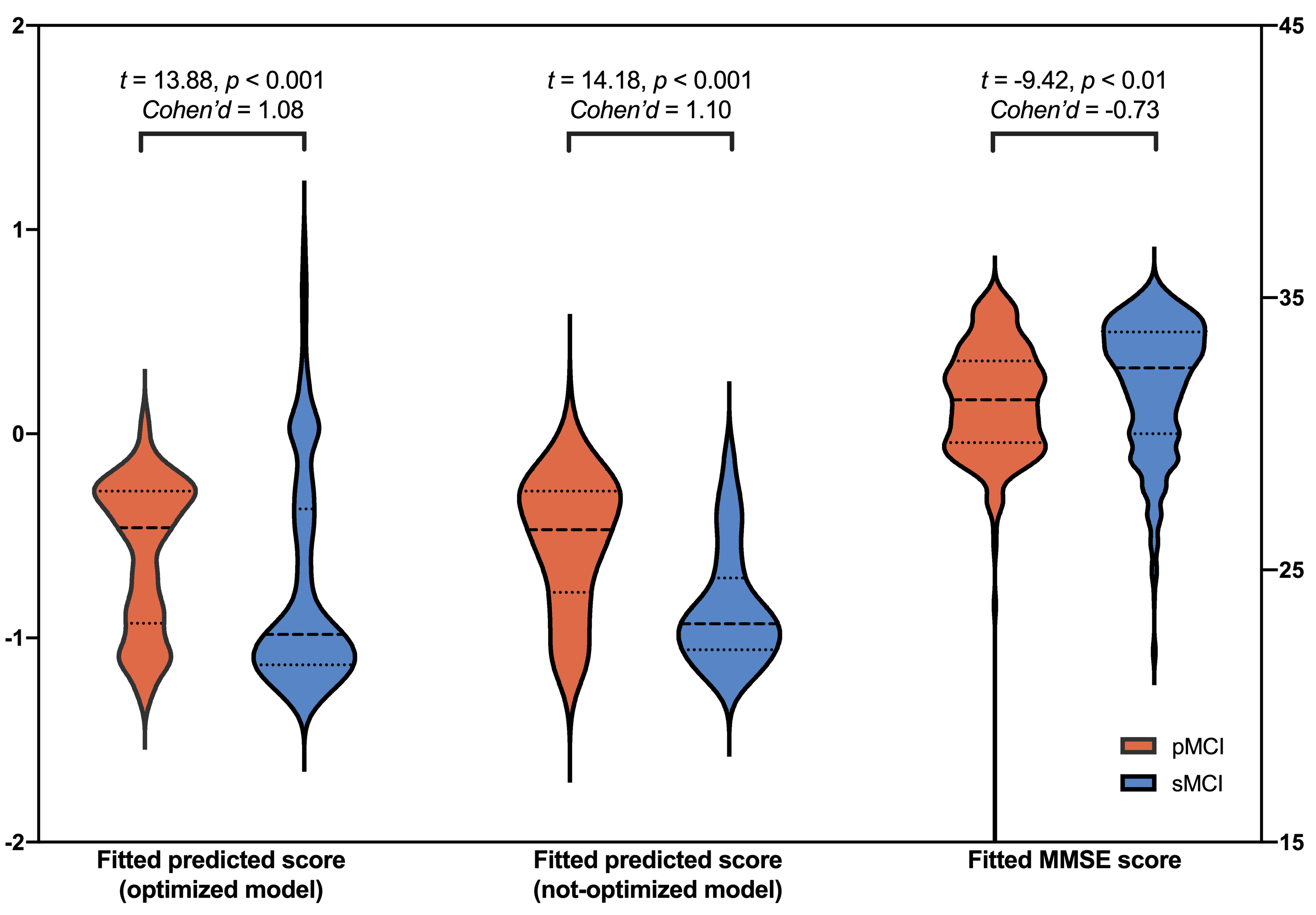

***Figure S8:* Violin plot of the predicted scores (output of the AD classifiers) and the mini-mental state examination (MMSE) scores of pMCI and sMCI participants in the first visit.** The predicted scores of AD classifiers and the MMSE scores were fitted values after regressing out age and sex.

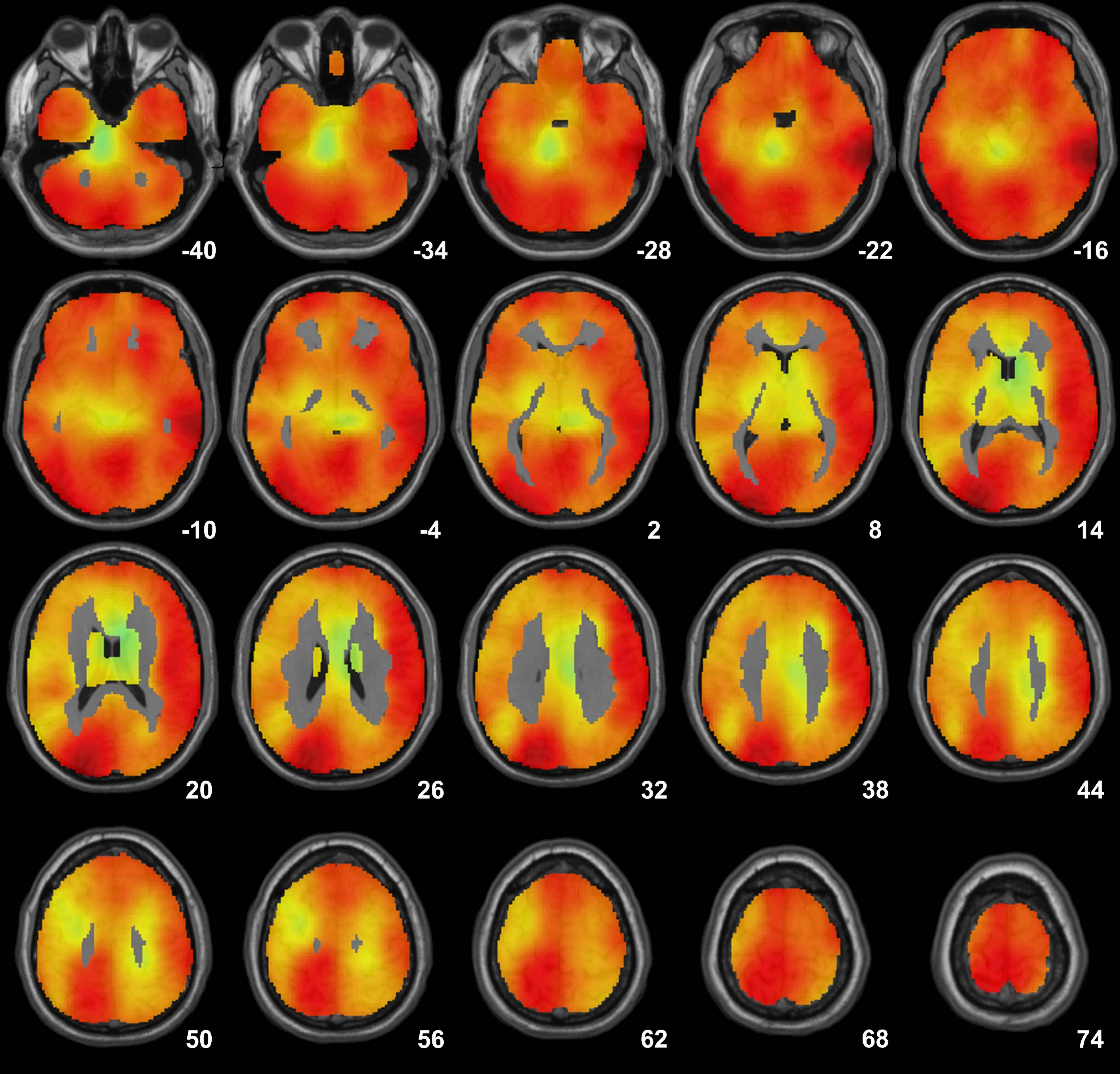

***Figure S9:* *Grad-CAM class activation map of participants in MIRIAD dataset.*** The Grad-CAM maps were generated for each sample and were averaged across all participants in the MIRIAD dataset, an independent testing dataset with high generalizability, to demonstrate where to attend for AD classification in an independent sample. Grad-CAM maps highlighted widespread grey matter areas, and the most important regions in occlusion maps also have relatively high weights in the Grad-CAM maps.

***Table S1:* Performance of the AD classifiers in the present study compared with some lightweight architectures**

|  | Accuracy | | | | AUC | | | | Sensitivity | | | | Specificity | | | |
| --- | --- | --- | --- | --- | --- | --- | --- | --- | --- | --- | --- | --- | --- | --- | --- | --- |
| Model structure | ADNI | AIBL | MIRIAD | OASIS | ADNI | AIBL | MIRIAD | OASIS | ADNI | AIBL | MIRIAD | OASIS | ADNI | AIBL | MIRIAD | OASIS |
| Inception-ResNet-V2 | 0.909  [0.902, 0.915] | 0.945  [0.926, 0.965] | 0.936  [0.917, 0.955] | 0.911  [0.895, 0.928] | 0.963  [0.958, 0.967] | 0.966  [0.945, 0.987] | 0.994  [0.991, 0.997] | 0.976  [0.966, 0.985] | 0.838  [0.824, 0.854] | 0.881  [0.816, 0.943] | 0.897  [0.869, 0.928] | 0.932  [0.889, 0.972] | 0.942  [0.935, 0.948] | 0.958  [0.941, 0.974] | 1.000  [1.000, 1.000] | 0.908  [0.890, 0.926] |
| DenseNet-201 | 0.823  [0.813, 0.832] | 0.822  [0.792, 0.853] | 0.960  [0.946, 0.978] | 0.681  [0.656, 0.708] | 0.936  [0.930, 0.942] | 0.942  [0.915, 0.969] | 0.996  [0.994, 0.998] | 0.962  [0.946, 0.977] | 0.913  [0.901, 0.925] | 0.890  [0.833, 0.947] | 0.990  [0.980, 1.000] | 0.985  [0.968, 1.000] | 0.933  [0.769, 0.795] | 0.809  [0.774, 0.842] | 0.911  [0.875, 0.949] | 0.639  [0.610, 0.669] |
| VGG19 | 0.870  [0.862, 0.878] | 0.897  [0.875, 0.920] | 0.921  [0.901, 0.942] | 0.837  [0.813, 0.867] | 0.927  [0.920, 0.933] | 0.946  [0.918, 0.974] | 0.977  [0.968, 0.985] | 0.952  [0.937, 0.967] | 0.782  [0.765, 0.780] | 0.871  [0.805, 0.941] | 0.874  [0.842, 0.905] | 0.932  [0.890, 0.972] | 0.911  [0.903, 0.919] | 0.902  [0.877, 0.927] | 1.000  [1.000, 1.000] | 0.824  [0.798, 0.846] |
| MobileNet-V2 | 0.816  [0.807, 0.825] | 0.871  [0.846, 0.897] | 0.916  [0.894, 0.939] | 0.803  [0.779, 0.826] | 0.907  [0.900, 0.914] | 0.924  [0.893, 0.954] | 0.979  [0.971, 0.988] | 0.920  [0.898, 0.942] | 0.840  [0.825, 0.855] | 0.83  [0.760, 0.905] | 0.933  [0.909, 0.958] | 0.836  [0.768, 0.900] | 0.805  [0.794, 0.817] | 0.878  [0.850, 0.905] | 0.889  [0.848, 0,927] | 0.798  [0.773, 0.822] |

The ADNI dataset contained 2,186 AD samples and 4,671 NC samples. The AIBL dataset contained 115 AD samples and 554 NC samples. The MIRIAD dataset contained 409 AD samples and 235 NC samples. The OASIS dataset contained 137 AD samples and 986 NC samples. The sample sizes shown here are the numbers of T1-weighted brain MRI scans. Abbreviations: AD = Alzheimer's disease participants, NC = normal control participants, GMD = grey matter density map, GMV = grey matter volume map, T1-weighted = normalized 3D T1-weighted structural image, AUC = area under the curve. The models listed here were derived from their 2D versions in keras-application and were modified for 3D MRI-derived feature map inputs and GPU capacity limits. The 95% confidence intervals are in brackets.
